# Automated Assignment of Proteoform Classification Levels

**DOI:** 10.1101/2021.05.18.444659

**Authors:** Zach Rolfs, Lloyd M. Smith

**Affiliations:** Department of Chemistry, University of Wisconsin-Madison, Madison, Wisconsin 53706

## Abstract

Proteoform identification is required to fully understand the biological diversity present in a sample. However, these identifications are often ambiguous because of the challenges in analyzing full length proteins by mass spectrometry. A five-level proteoform classification system was recently developed to delineate the ambiguity of proteoform identifications and to allow for comparisons across software platforms and acquisition methods. Widespread adoption of this system requires software tools to provide classification of the proteoform identifications. We describe here implementation of the five-level classification system in the software program MetaMorpheus, which provides both bottom-up and top-down identifications. Additionally, we developed a stand-alone program called ProteoformClassifier that allows users to classify proteoform results from any search program, provided that the program writes output that includes the information necessary to evaluate proteoform ambiguity. This stand-alone program includes a small test file and database to evaluate if a given program provides sufficient information to evaluate ambiguity. If the program does not, then ProteoformClassifier provides meaningful feedback to assist developers with implementing the classification system. We tested currently available top-down software programs and found that none of them other than MetaMorpheus provided sufficient information regarding identification ambiguity to permit classification.

## INTRODUCTION

Top-down proteomics is the premier method for identifying proteoforms, which are the various forms of proteins generated from a single gene.^1,2^ Despite recent advances in top-down proteomics, the identification of proteoforms remains difficult. Intact proteins frequently have poor fragmentation within the mass spectrometer, which makes the localization and identification of post-translational modifications (PTMs) nontrivial. In cases where the localization of a PTM is uncertain, search programs will either report multiple equally-probable proteoforms or arbitrarily pick a single proteoform to report. A deconvoluted intact mass could also be reported as a proteoform, despite the lack of any biological context. This uncertainty in the definition of a proteoform identification and the differences in how labs report results make comparisons across different software programs difficult.

A five-level proteoform classification system was previously introduced to address these issues caused by proteoform ambiguity.^3^ This system comprehensively considers sources of proteoform ambiguity arising from uncertain PTM localization, PTM identification, amino acid sequence, and gene of origin. Despite the clear need for this system in the proteoform community, adoption of the classification scheme has been limited to date to a single program, ProteoformSuite,^4^ which identifies proteoforms from intact mass measurements.^5^ More widespread adoption of this classification scheme is sorely needed in top-down proteomics.

We implemented the five-level proteoform classification system in the MetaMorpheus^6^ top-down search and provide methods for proteoform classification in the open-source software library mzLib (https://github.com/smith-chem-wisc/mzLib). The availability of this open-source code enables software developers to readily implement proteoform classification in both new and existing software programs. In an effort to expedite the community-wide adoption of proteoform classification, we additionally developed an open-source, standalone software program called ProteoformClassifier that can read generic proteoform identification results and classify each identified proteoform. Therefore, if an existing search program does not provide classification information, then ProteoformClassifier enables its users to categorize their proteoform output. Unfortunately, some programs are not transparent about ambiguity in their proteoform identifications. For example, if a PTM is equally likely to be on multiple amino acid residues but the search program only reports a PTM on a single residue, then that program is not transparent about ambiguity. Proteoform identifications from such programs cannot be accurately classified without changes to the existing program. We address this issue by providing a small test data file and database containing a proteoform from each of the possible classification levels. If a user is unsure whether or not their current top-down workflow is transparent about ambiguity, then they can analyze this test file with their workflow and validate the results with ProteoformClassifier. ProteoformClassifier checks that each proteoform result from the test file contains the expected ambiguity and provides helpful output if any unexpected issues are encountered.

## METHODS

### Classification Code

The previously published classification system focuses on four yes-or-no questions. For a given proteoform identification: 1) were the PTMs localized? 2) were the PTMs identified? 3) was the amino acid sequence identified? and 4) was the gene of origin identified? If all answers are “yes”, then the proteoform is a level 1 identification. If exactly three answers are “yes”, then it’s a level 2A, 2B, 2C, or 2D (depending on if the “no” answer is for question 1, 2, 3, or 4, respectively). If exactly two answers are “yes”, then it’s a level 3. If only one answer is “yes”, it’s a level 4. If all answers are “no”, then it’s a level 5.

The code for classifying proteoform identifications was written in C# and is freely available through mzLib. This code contains two methods that may be of interest to developers and both methods output a proteoform classification for a single identification. The first method accepts four Boolean values for each of the four possible sources of ambiguity discussed above. This method can be readily used in existing programs to provide classification information in their output and it is the method used by the MetaMorpheus top-down search. The second classification method accepts two text strings for the identification’s proteoform sequence(s) and gene(s). Multiple sequences or genes for a single identification are delimited with a vertical bar (e.g. ProteoformA|ProteoformB). PTMs are annotated with square brackets (e.g. [PTM and/or Mass]). This method parses the two text string inputs to determine ambiguity and then classifies the identification. The ability to read text strings enables the second method to parse generic input and is used by ProteoformClassifier.

### ProteoformClassifier

ProteoformClassifier is a free, open-source software tool that validates search program output and classifies proteoform identifications from validated search programs. It is available as a user-friendly graphical user interface and the command line at https://github.com/smith-chem-wisc/ProteoformClassifier/releases/. This program enables users to classify proteoform identifications from any search program by accepting a generic .tsv input file. The input for both validation and classification consists of three columns containing an arbitrary identifier (e.g. the scan number), the proteoform sequence(s), and the gene(s). Each row of the input is a unique proteoform-spectrum match (PrSM). A header is accepted, but not required. There are two output files produced by ProteoformClassifier when classifying proteoform identification results. The file “ClassifiedResults” contains the original three-column input with an additional fourth column for the classification level of each PrSM. The file “ClassifiedSummary” contains the number of PrSMs at each classification level, providing a clean summary of all results and allowing for facile comparisons across different workflows.

The validation module in ProteoformClassifier empowers users to determine if their current top-down search program is transparent about proteoform ambiguity. We created a small .mzML data file and .fasta protein database for users to test their bioinformatic workflow. This data file contains eight artificially constructed tandem mass spectra, each encoding a different proteoform with a different proteoform classification level *(Methods-Validation Data).* After analyzing this data file with a top-down search program, ProteoformClassifier can read the results and determine if all of the ambiguity that was expected was reported. If there is an issue with the output, it alerts the user that results from their workflow are not sufficiently transparent and cannot be accurately classified. In such cases, ProteoformClassifier additionally provides helpful output for developers to update their programs and implement the five-level proteoform classification system.

### Validation Data

We created a small top-down data file that contains eight tandem mass spectra, one for each of the possible eight proteoform classification levels (1, 2A, 2B, 2C, 2D, 3, 4, and 5). This data file was created by searching a previously published top-down yeast data file (05-26-17_B7A_yeast_td_fract7_rep1.raw)^7^ with MetaMorpheus^6^, TopPIC^8^, and MsPathFinder^9^. The eight tandem mass spectra came from PrSMs that were shared across all three search programs. All other tandem mass spectra (MS2) from the original file were removed. Precursor mass spectra (MS1) were retained if their scan number was within 100 scan numbers of a retained MS2 scan and their spectrum contained the precursor ion that was isolated for that retained MS2 scan. All other MS1 scans were removed. Retained MS1 spectra had their peaks filtered so that they only contained isotopic envelopes for the isolated proteoform of interest. This filtering step removed the possibility of MS1 deconvolution artifacts and ensured that the correct monoisotopic mass for each proteoform could be readily determined. The eight MS2 spectra had their fragment masses filtered such that only the b- and y-ions that matched to their proteoform identification remained. This filtering step was added because some top-down search programs consider additional fragment ions, such as neutral losses or internal fragments, which could be assigned and possibly used to disambiguate proteoform identifications. Removing these peaks enabled us to create intentionally ambiguous spectra and confidently generate a PrSM for each classification level. After filtering the MS2 fragment ions, we selected all MS2 spectra that encoded an ambiguous acetylated/trimethylated proteoform and added two new fragment ions. If the 42 Da mass shift associated with acetylation/trimethylation is unlocalized, it is often explained as an unambiguous N-terminal acetylation. We wanted to create PrSMs that had ambiguity in PTM identification, so we needed to localize the 42 Da mass shift to a lysine residue, where it could be explained by either an acetylation or a trimethylation. The two new fragment ions were added to both sides of the ambiguous PTM to ensure localization of the PTM to a lysine residue, rather than the proteoform N-terminus. Figure 1 shows how each of these eight proteoforms captures the ambiguity for their respective classification level. These PrSM results should be reproducible using any top-down software program when searching this test data file with the following settings: 5 ppm precursor and product mass tolerances, HCD (or file-specified) fragmentation, cleavage of initiator methionine, fixed carbamidomethylation (+57) of cysteine, variable oxidation of methionine, variable acetylation of lysine, and variable trimethylation of lysine.

**Figure 1.**
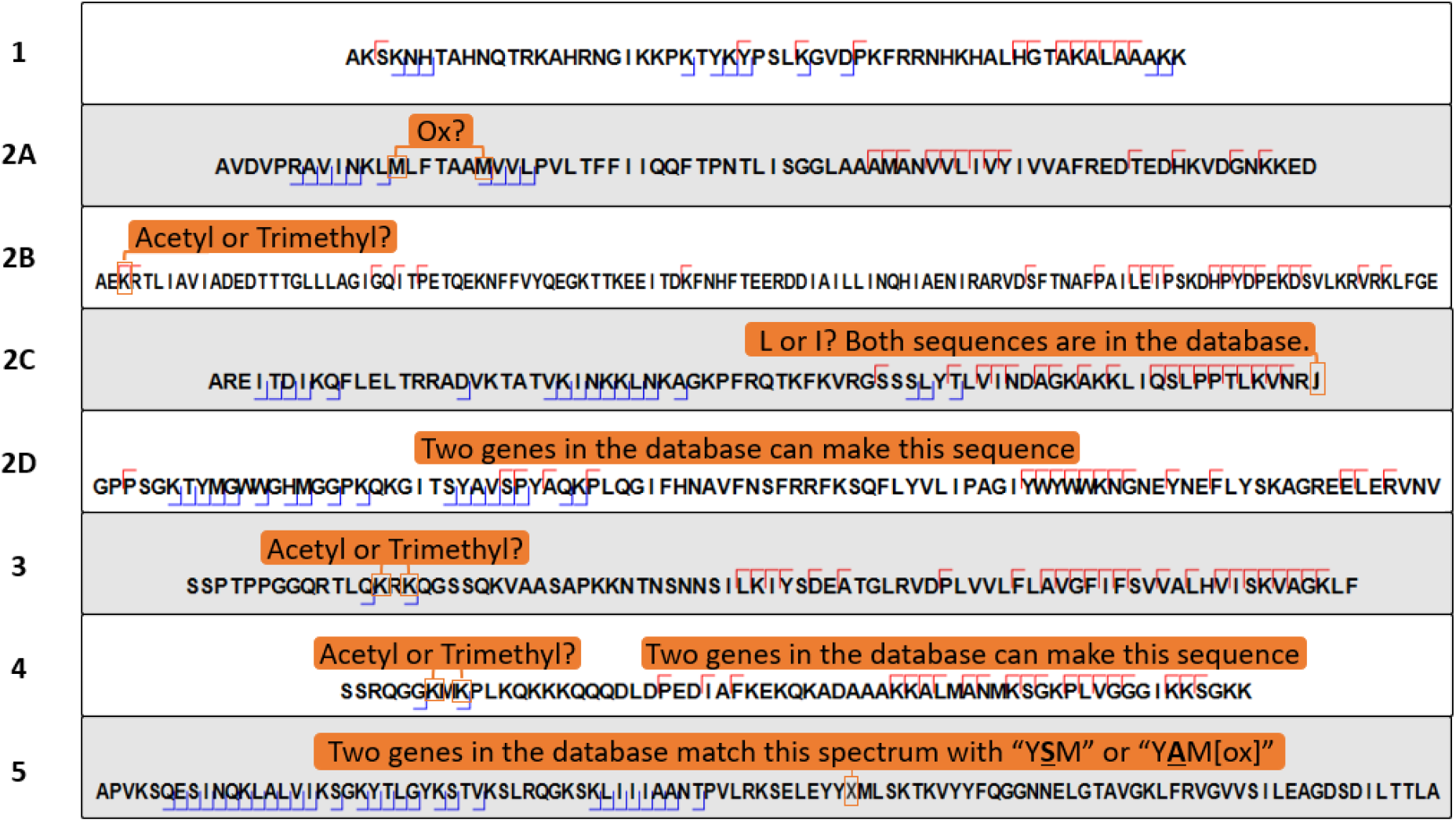
Proteoform-spectrum matches in the test data file and their corresponding classification levels (left side). The level 1 identification has all PTMs identified and localized with a single amino acid sequence from a single gene. Level 2A has an oxidation, but it’s uncertain if the oxidation is on the first or second methionine. Level 2B has a 42 Da shift localized to the first lysine, but the small mass difference between an acetylation and a trimethylation prevents a confident identification of the PTM. Level 2C is confidently identified, but there are two amino acid sequences originating from the same gene that score equally well. The leucine and isoleucine on the C-terminus cannot be disambiguated, making this a 2C. Level 2D is confidently identified, but there are two different genes in the database that produced the same protein sequence. This is a 2D because the gene of origin is uncertain. Level 3 has an unlocalized 42 Da shift, combining ambiguity from 2A and 2B. Level 4 has an unlocalized 42 Da shift and multiple genes can make this sequence (three sources of ambiguity from 2A, 2B, and 2D). Level 5 has all sources of ambiguity. Two different genes give rise to two different sequences that both explain the observed spectrum equally well. One of the sequences is unmodified, while the other has an oxidized methionine. Without knowing the gene, protein sequence, or PTMs, this last example is a level 5 identification.

## RESULTS

We evaluated ProteoformClassifier by searching the test data file (Methods-*Validation Data*) against MetaMorpheus^6^ v0.0.318, TopPIC^8^ v1.4.9, and MsPathFinder^9^ v1.1.7761. Other programs can be used but were not readily available during the writing of this manuscript. The test data file contains a proteoform spectrum for one of each proteoform classification level. These spectra were constructed to guarantee that the ambiguities could not be resolved. The test data were analyzed by all three programs with the results shown in Figure 2. MetaMorpheus provided proteoform results that were transparent about ambiguity from PTMs, amino acid sequences, and genes. TopPIC was transparent about ambiguity in PTM localization, but would arbitrarily report a single PTM, sequence, and gene when multiple equally probable options were available. There was a level 3 proteoform identified by TopPIC, which was caused because TopPIC reported a +42 Da mass shift rather than assigning an acetylation or trimethylation for that spectrum. An unidentified mass shift is automatically recognized by ProteoformClassifier and treated as an ambiguous PTM identification. MsPathFinder reported no ambiguity, despite the existence of ambiguity in seven of the eight proteoform spectra.

**Figure 2.**
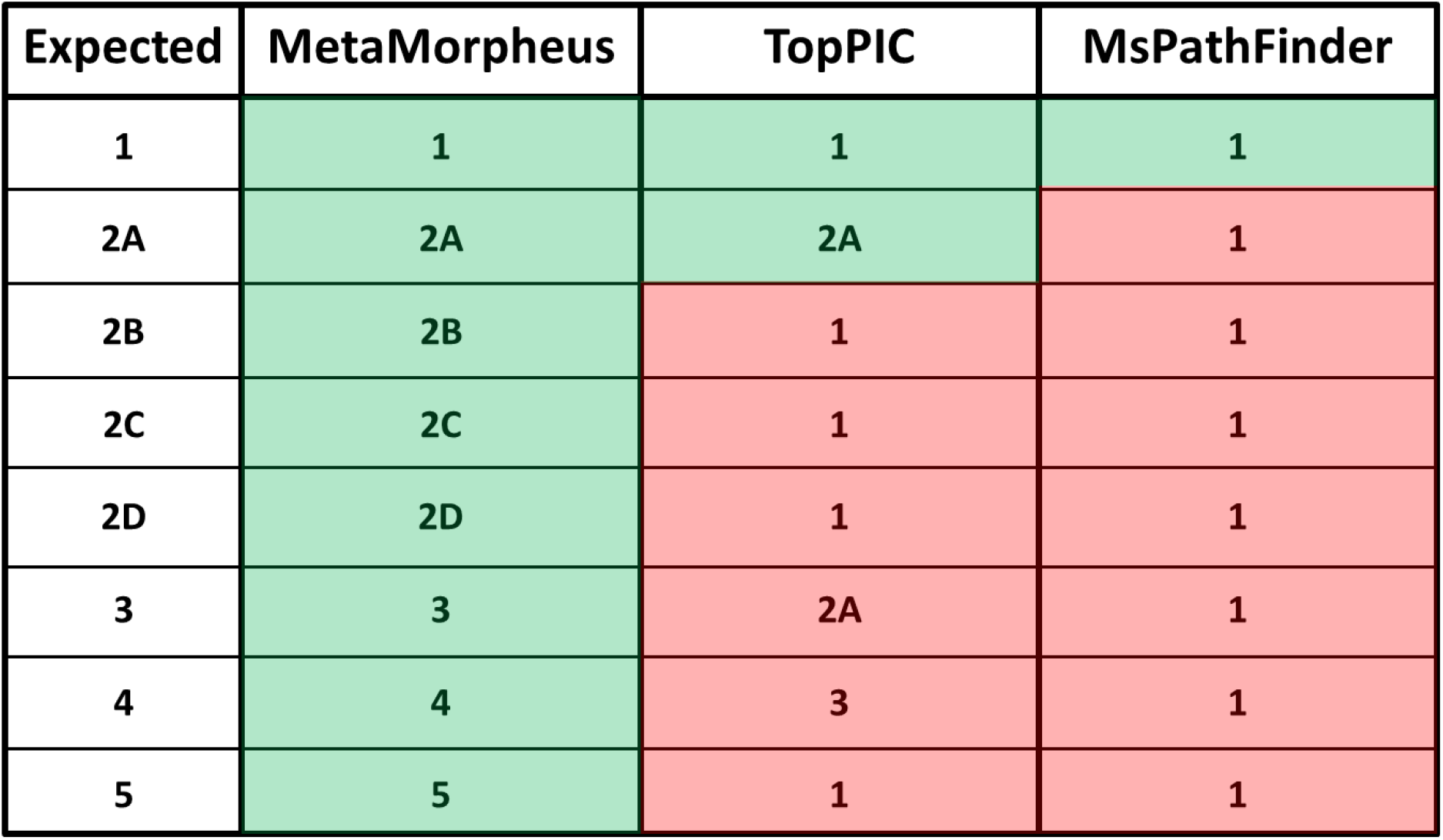
Classification results for each of the eight contrived spectra. Expected classifications for each spectrum are shown in the first column. The test file was analyzed by MetaMorpheus, TopPIC, and MsPathFinder. The results from these programs were provided as input for ProteoformClassifier and the classified results are shown in the three right columns. Green entries highlight output that correctly identified ambiguities. Red entries indicate that modifications are needed for the output to be accurately classified.

MetaMorpheus was the only available top-down search program that provided classifiable output. We analyzed a previously published yeast^7^ and human^10^ (Jurkat) dataset with MetaMorpheus to determine the extent of ambiguity in normal top-down search results. Mass calibration^6^ and Global-PTM-Discovery^11^ were used in MetaMorpheus prior to the final database search. A UniProtKB/Swiss-Prot.xml protein database of *Saccharomyces cerevisiae* (accessed Dec. 18, 2020) and *Homo sapiens* (accessed April 12, 2021) was used to search the two datasets. The default MetaMorpheus top-down settings were used for mass calibration, Global-PTM-Discovery, and the final database search.

The summarized classification results for these two datasets are shown in Table 1. The majority of PrSMs found at a 1% false-discovery rate were unambiguously identified, but over a third of PrSMs contained at least one source of ambiguity for both datasets. The most common source of ambiguity was PTM localization, which agrees with a previously classified intact mass dataset.^12^ Roughly 7-8% of PrSMs in both datasets had an ambiguous gene of origin. These PrSMs can be largely attributed to a few cases of genetic redundancy. For example, over 3,000 PrSMs in the Jurkat dataset came from an ATP synthase membrane subunit with ambiguity between ATP5G1, ATP5G2, and ATP5G3. The results for yeast were similar, where 490 PrSMs had gene ambiguity between 40S ribosomal protein S30 (RPS30A) and s30-B (RPS30B).

**Table 1.** Classification results for PrSMs identified at a 1% false-discovery rate for the yeast and Jurkat datasets.

| Classification | Yeast (# PrSMs) | Jurkat (# PrSMs) |
| --- | --- | --- |
| **1** | 9837 (67%) | 18252 (52%) |
| **2A** | 3040 (21%) | 10978 (31%) |
| **2B** | 17 (0%) | 264 (1%) |
| **2C** | 0 (0%) | 0 (0%) |
| **2D** | 1119 (8%) | 2585 (7%) |
| **3** | 512 (3%) | 2220 (6%) |
| **4** | 92 (1%) | 27 (0%) |
| **5** | 34 (0%) | 550 (2%) |
| **PrSMs with Ambiguity** | 4814 (33%) | 16624 (48%) |
| **Total PrSMs** | 14651 | 34876 |

Despite the low number of level 2B PrSMs, PTM identification ambiguity was observed in 83% of level 3 Jurkat PrSMs. The ambiguous PTMs were largely caused by uncertainty between acetylation and trimethylation, but also other PTMs such as methylation and oxidation. Despite the mass difference between methylation and oxidation, MetaMorpheus considers for missed monoisotopic mass shifts, allowing both PTMs to be assigned to the same spectrum without modifying the amino acid sequence. Unlike the Jurkat dataset, the yeast data found few instances of ambiguity in PTM identification. Only 9% of level 3 yeast PrSMs had an uncertain PTM identification, primarily between acetylation and trimethylation. Most level 3 yeast PrSMs had ambiguity in PTM localization (84%) and gene of origin (91%). These two sources of ambiguity were also sizeable for level 3 Jurkat PrSMs (83% for PTM localization and 34% for gene of origin).

## DISCUSSION

The five-level proteoform classification system is an important contribution to the top-down community. This system defines what a proteoform identification means and enables proteoform ambiguity to be better communicated and compared across software programs. Currently, the system has yet to be widely adopted by the community. With the exception of MetaMorpheus, none of the top-down software programs that we tested were transparent about their ambiguity. Understanding the uncertainty in an identification is important for researchers who are attempting to adopt new methods or replicate published studies. The lack of transparency in existing software tools remains a major obstacle in the community-wide adoption of the classification system and consequently inhibits progress in the field.

We developed the open-source program ProteoformClassifier to aid developers in the implementation of the classification system and provide users with a means to classify their proteoform results. This software program can accurately classify proteoform identifications that are transparent about ambiguity in PTM localization, PTM identification, proteoform sequence, and gene of origin. ProteoformClassifier additionally provides helpful feedback to elucidate which ambiguities, if any, are not currently reflected in a given search program’s output through the aid of a contrived dataset with known ambiguities. Finally, the classification code that we created has been made available in the software library mzLib, allowing developers to easily implement the classification method once their software is transparent about proteoform ambiguity.

## AUTHOR INFORMATION

*Corresponding Author: Phone: (608) 263-2594.

## Author Contributions

Z.R. developed ProteoformClassifier and wrote the manuscript. L.M.S. edited the manuscript and directed the project. The authors declare no competing financial interest.

## ACKNOWLEDGEMENTS

This work was supported by the NIH National Institute of General Medical Sciences under award number R35GM126914. The content is solely the responsibility of the authors and does not necessarily represent the official views of the NIH. We thank Michael R. Shortreed, Katherine E. Henke, and Kyndalanne Pike for their help in testing ProteoformClassifier.

## ABBREVIATIONS

PTM: post-translational modification;
PrSM: proteoform-spectrum match;
MS1: precursor mass spectrum;
MS2: tandem mass spectrum.

## REFERENCES

(1) Smith, L.M.; Kelleher, N.L.; Consortium for Top Down Proteomics. Proteoform: a single term describing protein complexity. Nat. Methods 2013, 10, 186–187.

(2) Smith, L.M.; Kelleher, N.L. Proteoforms as the next proteomics currency. Science 2018, 359(6380), 1106–1107.

(3) Smith, L.M.; Thomas, P.M.; Shortreed, M.R.; Schaffer, L.V.; Fellers, R.T.; LeDuc, R.D.; Tucholski, T.; Ge, Y.; Agar, J.N.; Anderson, L.C.; Chamot-Rooke, J.; Gault, J.; Loo, J.A.; Pasa-Tolic, L.; Robinson, C.V.; Schluter, H.; Tsybin, Y.O.; Vilaseca, M.; Vizcaino, J.A.; Danis, P.O.; Kelleher, N.L. A five-level classification system for proteoform identifications. Nat. Methods 2019 16, 939–940 (2019).

(4) Cesnik, A.J.; Shortreed, M.R.; Schaffer, L.V.; Knoener, R.A.; Frey, B.L.; Scalf, M.; Solntsev, S.K.; Dai, Y.; Gasch, A.P.; Smith, L.M. Proteoform Suite: Software for Constructing, Quantifying, and Visualizing Proteoform Families. J. Proteome Res. 2018, 17 (1), 568–578.

(5) Shortreed, M.R.; Frey, B.L.; Scalf, M.; Knoener, R.A.; Cesnik, A.J.; Smith, L.M. Elucidating Proteoform Families from Proteoform Intact-Mass and Lysine-Count Measurements. J. Proteome Res. 2016, 15 (4), 1213–1221.

(6) Solntsev, S.K.; Shortreed, M.R.; Frey, B.L.; Smith, L.M. Enhanced Global Post-Translational Modification Discovery with MetaMorpheus. J. Proteome Res. 2018, 17 (5), 1844–1851.

(7) Schaffer, L.V.; Shortreed, M.R.; Cesnik, A.J.; Frey, B.L.; Solntsev, S.K.; Scalf, M.; Smith, L.M. Expanding Proteoform Identifications in Top-Down Proteomic Analyses by Constructing Proteoform Families. Anal. Chem. 2018, 90 (2), 1325–1333.

(8) Kou, Q.; Xun, L.; Liu, X. TopPIC: a software tool for top-down mass spectrometry-based proteoform identification and characterization. Bioinformatics 2016, 32 (22), 3495–3497.

(9) Park, J.; Piehowski, P.D.; Wilkins, C.; Zhou, M.; Mendoza, J.; Fujimoto, G.M.; Gibbons, B.C.; Shaw, J.B.; Shen, Y.; Shukla, A.K.; Moore, R.J.; Liu, T.; Petyuk, V.A.; Tolic, N.; Pasa-Tolic, L.; Smith, R.D.; Payne, S.H.; Kim, S. Informed-Proteomics: open-source software package for top-down proteomics. Nat. Methods 2017, 14, 909–914.

(10) Dai, Y.; Buxton, K.E.; Schaffer, L.V.; Miller, R.M.; Millikin, R.J.; Scalf, M.; Frey, B.L.; Shortreed, M.R.; Smith, L.M. Constructing Human Proteoform Families Using Intact-Mass and Top-Down Proteomics with a Multi-Protease Global Post-Translational Modification Discovery Database. J. Proteome Res. 2019, 18, 3671–3680

(11) Li, Q.; Shortreed, M.R.; Wenger, C.D.; Frey, B.L.; Schaffer, L.V.; Scalf, M.; Smith, L.M. Global Post-Translational Modification Discovery. J. Proteome Res. 2017, 16 (4), 1383–1390.

(12) Schaffer, L.V.; Millikin, R.J.; Shortreed, M.R.; Scalf, M.; Smith, L.M. Improving Proteoform Identifications in Complex Systems Through Integration of Bottom-Up and Top-Down Data. J. Proteome Res. 2020, 19 (8), 3510–3517.

